# Stair Recognition for Robotic Exoskeleton Control using Computer Vision and Deep Learning

**DOI:** 10.1101/2022.04.11.487925

**Authors:** Andrew Garrett Kurbis, Brokoslaw Laschowski, Alex Mihailidis

**Affiliations:** Biomedical Engineering Program, University of Waterloo, ON Canada, on placement in the Temerty Faculty of Medicine, University of Toronto, ON Canada; Temerty Faculty of Medicine, University of Toronto, ON Canada, and the Toronto Rehabilitation Institute, ON Canada

## Abstract

Computer vision can be used in robotic exoskeleton control to improve transitions between different locomotion modes through the prediction of future environmental states. Here we present the development of a large-scale automated stair recognition system powered by convolutional neural networks to recognize indoor and outdoor real-world stair environments. Building on the *ExoNet* database - the largest and most diverse open-source dataset of wearable camera images of walking environments – we designed a new computer vision dataset, called *StairNet*, specifically for stair recognition with over 515,000 images. We then developed and optimized an efficient deep learning model for automatic feature engineering and image classification. Our system was able to accurately predict complex stair environments with 98.4% classification accuracy. These promising results present an opportunity to increase the autonomy and safety of human-exoskeleton locomotion for real-world community mobility. Future work will explore the mobile deployment of our automated stair recognition system for onboard real-time inference.

## I. Introduction

Human locomotion follows a biological system feedback loop [1], [2], which can be described by four main processes: 1) recognition of the physical environment using the human visual system; 2) cognitive processing of the environmental states and locomotor intent via neural control; 3) translation of the locomotor intent to movement through the musculoskeletal (MSK) system; and 4) physical environment response (i.e., the new walking environment that the human interacts with). However, if this feedback loop is disrupted by limitations to the musculoskeletal system due to aging and/or physical disabilities such as osteoarthritis, or communication limitations to the central nervous system due to stroke or spinal cord injury, this can affect one’s ability to perform daily locomotor activities and navigate new challenging environments safely and effectively [3].

Robotic lower-limb exoskeletons may help address these limitations and allow users to regain mobility and independence through powered locomotor assistance via motorized joints [4]. Similar to the biological vision-locomotor feedback loop, automated high-level control of human-exoskeleton locomotion for real-world mobility requires continuous assessment of future environmental states for seamless transitions between different locomotion modes, each of which include individual control parameters. Accurate classification of stair environments can be particularly important due to the safety implications and the greater theoretical risk of serious injuries if the environment is misclassified during stair ascent.

Although stair recognition systems have been developed for wearable robotics [5]–[13], especially powered prosthetic legs, these systems have mainly been limited to statistical pattern recognition and machine learning algorithms that require manual feature engineering and/or have involved relatively small image datasets, which can restrict their real-world application and generalizability to complex environments [14]. Motivated by these limitations, here we present the development of a large-scale automated stair recognition system powered by computer vision and deep learning to recognize indoor and outdoor real-world stair environments (Figure 1). The longterm objective of this research is to develop next-generation environment-adaptive control systems for robotic exoskeletons and other mobility assistive technologies.

**Figure 1.**
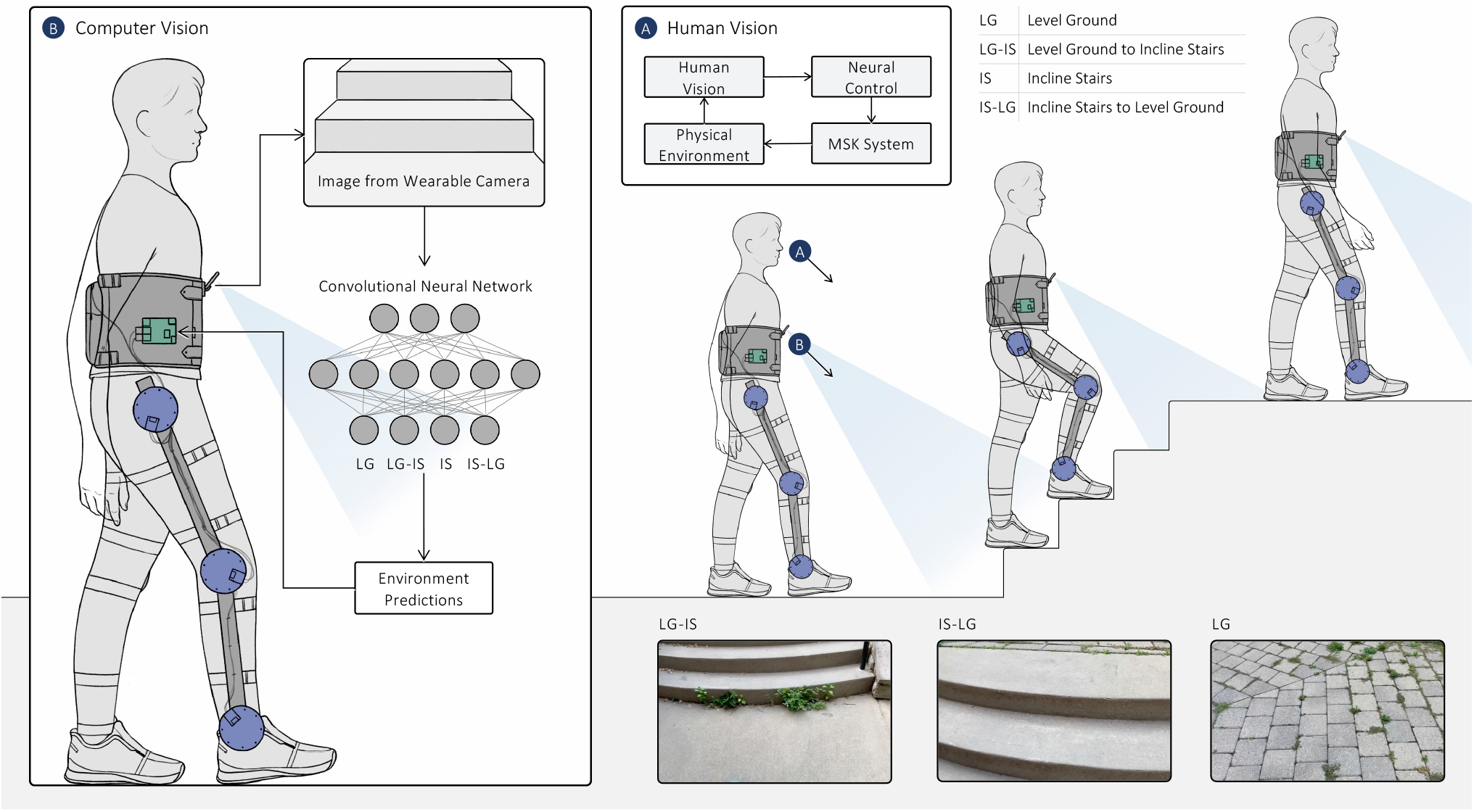
Schematic showing the application of computer vision and deep learning for sensing and classification of real-world stair environments during human-exoskeleton locomotion.

## II. Methods

### A. Image Dataset

A customized image dataset was created using the manually annotated videos from the *ExoNet* database [15] – the largest and most diverse open-source dataset of wearable camera images of walking environments. Images from six of the twelve original ExoNet classes were used, which included environments that an exoskeleton user would encounter when ascending stairs. The initial dataset included ~543,000 RGB images. The “level ground” and “incline stairs” steady-state classes in the ExoNet database included separate classes for images that contained doors/walls and those that did not. This differentiation was not applied to our study such that the six ExoNet classes were combined into four. The final four environment classes in our dataset included: level ground (LG), which included both ExoNet classes “level ground steady state” and “level ground transition to door/wall”; level ground – incline stairs (LG-IS), which consisted of the ExoNet class “level ground transition to incline stairs”; incline stairs (IS), which included both ExoNet classes “incline stairs steady state” and “incline stairs transition to door/wall”; and incline stairs – level ground (LG-IS), which consisted of the ExoNet class “incline stairs transition to level ground.” The dataset was randomly split into training (89.5%), validation (3.5%), and testing (7%) sets, matching the subset distribution values from Laschowski and colleagues [16], while maintaining the class distributions– i.e., 85.8% for LG, 9.3% for IS, 3.1% for LG-IS, and 1.8% for IS-LG.

### B. Convolutional Neural Network

A convolutional neural network (CNN) model was developed using the base model of MobileNetV2 [17], [18] (Table 1). MobileNetV2 is a CNN architecture based on depthwise separable convolutions. The model uses width and resolution multipliers to create a lightweight framework that trades a small amount of accuracy for reduced computational requirements. The model was chosen for its efficient and lightweight design, allowing for future consideration for onboard computations and robotic exoskeleton control. The MobileNetV2-based model was created in TensorFlow 2.7 [19]. Different model parameters were tested using a Google Cloud Tensor Processing Unit (TPU), a hardware accelerator optimized for efficient machine learning through large matrix operations, in order to optimize the system performance. An initial model was developed using the parameter ranges from Laschowski and colleagues [20], [21], who trained MobileNetV2 and more than a dozen other deep learning models on the entire ExoNet database. Our initial model design included the base model of MobileNetV2; the Adam optimizer [22]; four classes with randomly initialized weights; ~2.3 million parameters; a base learning rate of 0.001; a batch size of 128; and a cosine weight decay learning rate policy.

**Table 1.** The design of the MobileNetV2 convolutional neural network [18], which was used for automatic feature engineering and image classification of real-world stair environments. The network has ~2.3 million total parameters.

| Network Stage | Operator | Input Resolution | Output Channels | Number of Layers | Number of Parameters |
| --- | --- | --- | --- | --- | --- |
| 1 | Conv2d | 224x224x3 | 32 | 1 | 992 |
| 2 | Bottleneck | 112x112x32 | 16 | 1 | 932 |
| 3 | Bottleneck | 112x112x16 | 24 | 6 | 5,568 |
| 4 | Bottleneck | 56x56x24 | 32 | 6 | 20,096 |
| 5 | Bottleneck | 28x28x32 | 64 | 6 | 53,372 |
| 6 | Bottleneck | 14x14x64 | 96 | 6 | 236,160 |
| 7 | Bottleneck | 14x14x96 | 160 | 6 | 399,424 |
| 8 | Bottleneck | 7x7x160 | 320 | 6 | 1,126,720 |
| 9 | Conv2d | 7x7x320 | 1280 | 1 | 414,720 |
| 10 | Avgpool 7x7 | 7x7x1280 | 1 | 1 | 0 |
| 11 | Dropout | 1280 | 1 | 1 | 0 |
| 12 | Dense | 1280 | 1 | 1 | 5124 |

### C. Performance Evaluation

We used the following metrics to quantitatively evaluate the model performance: accuracy *(A), f1* score, precision *(P),* and recall value *(R)*.

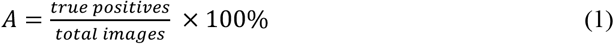

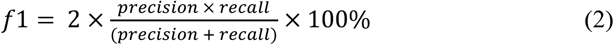

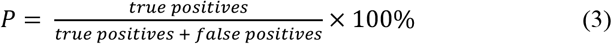

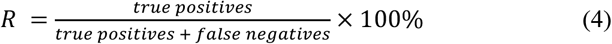

### D. StairNet Dataset

Initial experimentation was performed on the validation set using a preliminary optimized CNN model. Low categorical accuracies of 53.6% and 66.4% were found for the transition classes IS-LG and LG-IS, respectively. An investigation into the cause of low accuracy revealed a number of misclassified images and images with significant obstructions in the ExoNet database. The number of images with significant obstructions were most abundant in the LG class. We manually resorted the images and created a new dataset named *StairNet*. This dataset was developed to be used for future research in automated stair recognition and to prevent the misclassification of environments in real-world applications due to automatic feature extraction of misclassified images or images with significant obstructions presented during model training. Table 2 outlines the new class definitions that were developed to outline a more precise cut-off point between the different environment classes when re-sorting the images. After three manual annotation passes of the dataset, images that were considered out of scope (e.g., an image of a wall without level ground visible) and images that had significant obstructions were discarded, therein reducing the total number of images in our new dataset to ~515,000. After completing the dataset revision, the deep learning model was retested on the validation set with the same parameters as previously used. The low categorical accuracies improved from 53.6% to 84.4% for the IS-LG class and from 66.4% to 87.5% for the LG-IS class. We then randomly split the dataset into training (89.5%), validation (3.5%), and testing (7%) sets while maintaining the class distributions (Table 3). The StairNet dataset was uploaded to IEEE DataPort and is available for download https://ieee-dataport.org/documents/stairnet-computer-vision-dataset-stair-recognition [23].

**Table 2.** Description of the environment classes, which were used to manually label our new computer vision dataset called StairNet.

| Class | Image Description |
| --- | --- |
| LG | An image that contains a level ground environment where incline stairs are not clearly visible. |
| LG-IS | An image with incline stairs where the horizontal surface area of the bottom step or landing is clearly greater than the surface area of other steps apparent in the image (i.e., the surface area or depth is approximately 1.5x the size of subsequent steps). |
| IS | An image with multiple incline stairs where the horizontal surface area of the top and bottom step or landing are not clearly greater than one another. |
| IS-LG | An image with incline stairs where the horizontal surface area of the top step or landing is clearly greater than that of other steps or landings apparent in the image (i.e., the surface area or depth is approximately 1.5x the size of subsequent steps). For an incline stair to be included in the IS-LG class, the horizontal face of the last step prior to the level ground must be visible. |

**Table 3.**
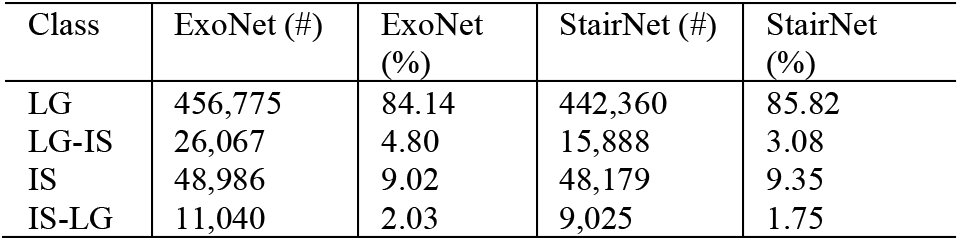
Distribution of the environment classes in the ExoNet database [15] and our new computer vision dataset – StairNet. The number of images (#) and the percent of the dataset (%) are reported for each class.

### E. Model Optimization

The model parameters were tuned with the StairNet dataset and tested using different model designs to optimize the validation accuracy and loss. We first developed a model using transfer learning with pre-trained weights from *ImageNet* [24]. We also added a global average pooling 2D layer and a softmax dense prediction layer, which decreased the number of trainable parameters, thus reducing the time and computational requirements to optimize the model. Five freeze layer combinations were tested to evaluate the use of transfer learning at 141, 100, 50, 25, and 5 frozen layers. The different combinations were run for 60 epochs. The optimal freeze layer combination for the model was 5 frozen layers with ~2.2 million parameters, which resulted in the highest validation accuracy and lowest validation loss.

We then optimized the model for different batch sizes (64, 128, 256) and base learning rates (0.0001, 0.00001, 0.000001). Each model design was run for 60 epochs. The optimal combination was a batch size of 256 and a base learning rate of 0.0001. We then compared the optimized pre-trained model to the equivalent non-pretrained model with randomly initialized weights. After 60 epochs, both models started to plateau. The pre-trained model had a higher overall validation accuracy (~98%) compared to the non-pre-trained model (~97%) while having similar final validation loss values. After 60 epochs, the pre-trained model exhibited characteristics of overfitting such that an increase in the validation loss was observed as the models approached 60 epochs. We compared the optimized pre-trained model to variations with an additional dropout layer. The model variations included dropout rates of 0.01, 0.02, and 0.05. The additional dropout layer with a dropout rate of 0.02 resulted in the highest validation accuracy and lowest validation loss. We then compared the previous model to a variation that used L2 weight regularization. The models were run for 60 epochs, and the validation loss and accuracy were compared, which showed no additional performance benefit of weight regularization.

We also explored oversampling of underrepresented classes (e.g., IS-LG and LG-IS) to reduce overfitting. The model was oversampled by randomly resampling and augmenting images previously presented to the model during training. Five model designs were compared, which included different values for the minimum number of images per class required for model training – i.e., 25,000; 40,000; 60,000; 200,000; and 400,000. There was a decrease in the overall validation accuracy as the minimum value increased. However, the categorical accuracy for the underrepresented classes increased as the minimum value increased, thereby creating a more even categorical accuracy throughout the different classes (Tables 4 and 5). Since there could be more significant consequences resulting from a false negative than a false positive in the detection of stairs for robotic exoskeleton control, we decided to select the deep learning model that oversampled with a minimum value of 400,000 images per class since this model had an even categorical accuracy distribution and the lowest probability of false negatives as seen in the reduced probability of misclassification as LG in class IS (0.3%) and IS-LG (2.2%).

**Table 4.**
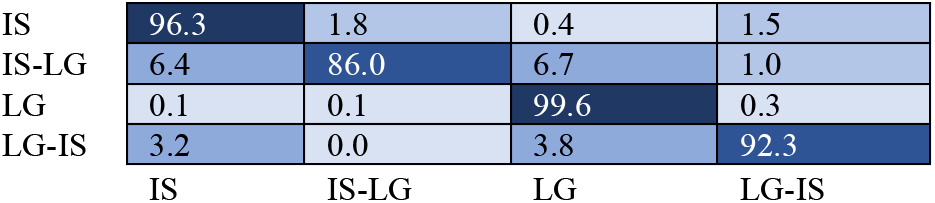
Normalized confusion matrix of the environment predictions for the validation set using the oversampled model with a minimum value of 25,000 images per class. The horizontal and vertical axes are the predicted and labelled classes, respectively.

**Table 5.** Normalized confusion matrix of the environment predictions for the validation set using the oversampled model with a minimum value of 400,000 images per class. The horizontal and vertical axes are the predicted and labelled classes, respectively.

|  |  |  |  |  |
| --- | --- | --- | --- | --- |
| IS | 96.6 | 1.3 | 0.3 | 1.8 |
| IS-LG | 5.7 | 90.8 | 1.9 | 1.6 |
| LG | 0.1 | 0.3 | 99.0 | 0.6 |
| LG-IS | 3.2 | 0.4 | 2.2 | 94.2 |
|  | IS | IS-LG | LG | LG-IS |

We then fine-tuned the oversampled model with different batch sizes and base learning rates on a high epoch run to further increase the categorical and overall validation accuracies. By comparing the validation accuracy and loss as well as the confusion matrices, the final model design was selected with a reduced base learning rate of 0.00001, a batch size of 128, and a cosine weight decay learning policy. The final MobileNetV2-based model included the use of pretrained weights, 5 frozen layers, and ~2.2 million parameters. The model was oversampled with a minimum categorical image count of 400,000 images and contained an additional dropout layer with a dropout rate of 0.02. The final model was run for 60 epochs (Figure 2).

**Figure 2.**
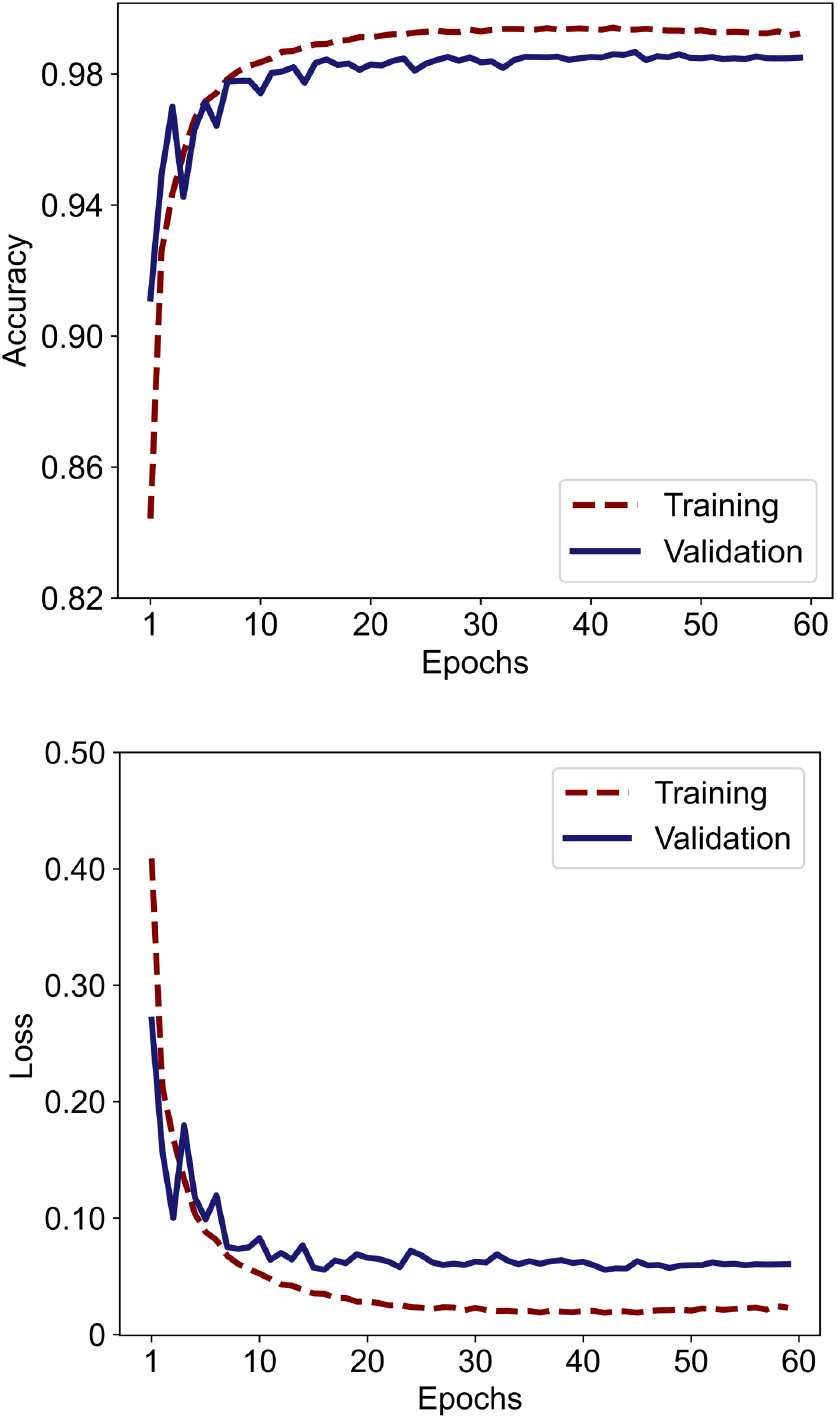
The image classification accuracy and loss for the training and validation sets over 60 epochs with the optimized deep learning model parameters.

## III. Results

The image classification accuracies on the training and validation sets were 99.3% and 98.5%, respectively. When evaluated on the testing set, the deep learning model achieved an overall classification accuracy *(A)* of 98.4%, a weighted *f1* score of 98.4%, a weighted precision value *(P)* of 98.5%, and a weighted recall value *(R)* of 98.4%. Here, the classification accuracy is defined as the number of true positives identified by the neural network (35,507 images) out of the total number of images in the testing set (36,085).

The image classification accuracy on the testing set varied between the different environment classes with a categorical accuracy of 99.0% for LG, 91.7% for LG-IS, 96.9% for IS, and 90.5% for IS-LG. Table 6 shows the normalized confusion matrix, which illustrates the classification performance distribution during inference. The two transition classes (i.e., LG-IS and IS-LG) achieved the lowest categorical accuracies, which can likely be attributed to having the smallest class distributions in the dataset, comprising only ~3.1% and ~1.8% of the total number of images, respectively.

**Table 6.** Normalized confusion matrix of the environment predictions for the testing set. The horizontal and vertical axes are the predicted and labelled classes, respectively.

|  |  |  |  |  |
| --- | --- | --- | --- | --- |
| IS | 96.9 | 1.5 | 0.3 | 1.3 |
| IS-LG | 6.5 | 90.5 | 2.4 | 0.6 |
| LG | 0.1 | 0.3 | 99.0 | 0.7 |
| LG-IS | 4.3 | 0.5 | 3.4 | 91.7 |
|  | IS | IS-LG | LG | LG-IS |

Figure 3 shows several examples of failure cases. The first row contains images from the LG class that were incorrectly classified as LG-IS. The images contain features common within the LG-IS class such as strong horizontal lines in the top section of the images. These horizontal line features were likely extracted by the neural network and confused as transitions to incline stairs. The second row contains LG images that were incorrectly classified as IS. The images contain strong horizontal lines resulting from surface textures (e.g., brick flooring in the second column). These horizontal line features are present throughout the image and were likely confused for steady-state incline stairs. The bottom row contains images from the stair classes that were misclassified as LG; these misclassifications pose the most significant safety risk to exoskeleton users. The images contain unique stair characteristics like unusual materials (e.g., wood) or angles, which rotate the horizontal features to a diagonal or vertical axis.

**Figure 3.**
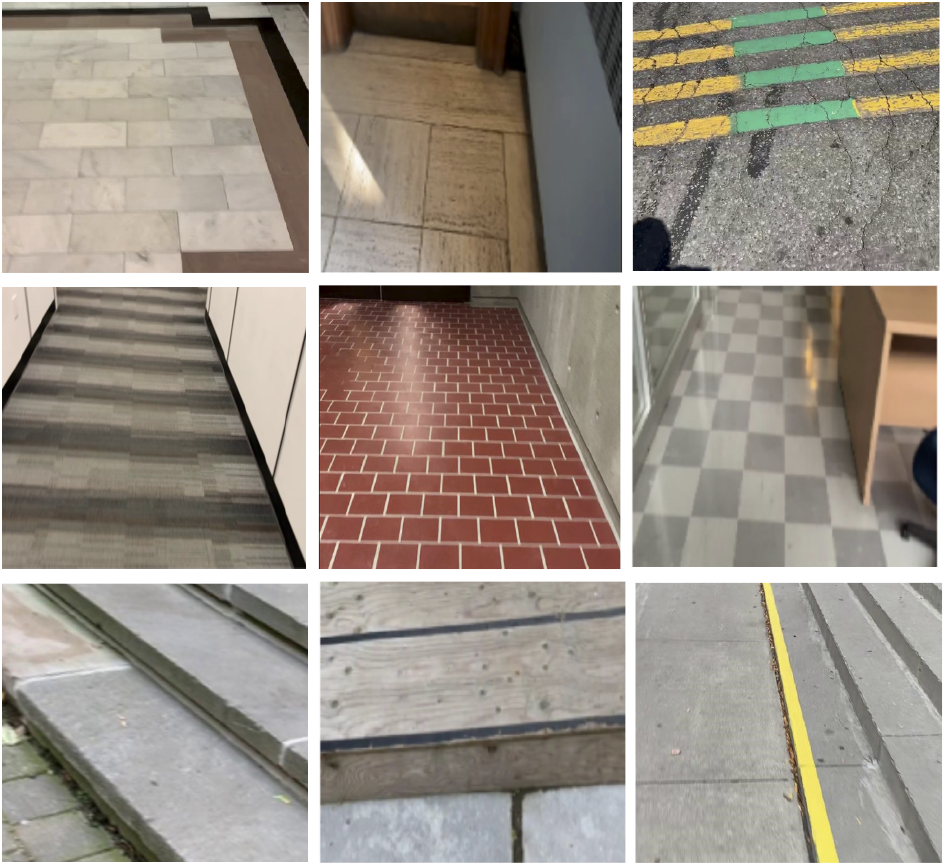
Examples of incorrect environment predictions by the convolutional neural network. The first row contains level ground images that were misclassified as level ground – incline stairs (LG-IS). The second row contains level ground images that were misclassified as incline stairs (IS). The third row contains images from the stair classes that were misclassified as level ground (LG).

## IV. Discussion

Here we developed a large-scale automated stair recognition system powered by computer vision and deep convolutional neural networks to recognize indoor and outdoor real-world stair environments. These classification predictions could be used to develop environment-adaptive control systems for robotic exoskeletons and other mobility assistive technologies. Building on the ExoNet database [15] – the largest and most diverse open-source dataset of wearable camera images of walking environments – we designed a new computer vision dataset named StairNet, specifically for stair recognition. We then optimized an efficient deep learning model [17], [18] for automatic feature engineering and image classification. Our stair recognition system was able to accurately predict complex stair environments with 98.4% overall classification accuracy. These promising results present an opportunity to increase the autonomy and safety of human-exoskeleton locomotion for real-world community mobility.

Compared to the classification results for the stair classes in the original ExoNet database [16], [20], [21], we achieved significantly higher prediction accuracies (Appendix 1). First, we discovered a number of ambiguous labelled images in the ExoNet classes LG-S, LG-T-DW, LG-T-IS, IS-S, IS-T-DW, and IS-T-LG. We manually re-sorted these images into four classes (i.e., LG, LG-IS, IS, and IS-LG) using new definitions to increase the precision of the cut-off points between class transitions. Prior to the dataset revision, a preliminary stair recognition model was developed, which achieved validation accuracies of 53.6% and 66.4% for the transition classes IS-LG and LG-IS, respectively; these results are similar to previous research [20]. We then retested the same model on the StairNet dataset and the prediction accuracies significantly increased to 84.4% for IS-LG and 87.5% for LG-IS. Our new computer vision dataset for stair recognition contains ~515,000 images distributed across four classes.

Compared to previous stair recognition systems for wearable robotics [5]–[13], our classification system has several advantages (Appendix 1). Many researchers have used statistical pattern recognition and machine learning algorithms that require time-consuming and suboptimal hand engineering. In contrast, our deep learning model replaces hand-designed features with multilayer networks that can automatically and efficiently learn the optimal image features from training data. Furthermore, compared to other CNN-based stair recognition systems, our system is significantly larger. For instance, Laschowski and colleagues [6] developed one of the first automated stair recognition systems using convolutional neural networks. However, their dataset included only ~34,000 labelled images. In comparison, StairNet includes over 515,000 labelled images. These differences can have important practical implications since deep learning often requires significant and diverse training data to facilitate generalization to complex real-world applications [14].

Despite these advances, our study has several limitations. We used a high-performance Google Cloud TPU for training and testing our deep learning model. This additional computing power supported the large-scale hyperparameter optimization and the resulting predictive capabilities of the model design. However, it is relatively unknown how the classification performance on these AI accelerator application-specific integrated circuits will translate to mobile and embedded computing devices for robotic exoskeletons. Although we used an efficient, lightweight convolutional neural network [17], [18] with low computational requirements to increase the potential for onboard real-time inference, the feasibility of model deployment was not evaluated in this study. Further investigation is needed to assess the computing performance of this lightweight CNN model on mobile devices. We plan to address these limitations in future work.

Overall, the promising results of our automated stair recognition system using computer vision and deep learning demonstrate the potential for applications in robotic exoskeleton control to increase user safety and autonomy during locomotion through the highly accurate prediction of future environmental states.

## Acknowledgments

We want to recognize William McNally, Alexander Wong, and John McPhee from the University of Waterloo for their efforts on the original ExoNet database. We also acknowledge Shehryar Saharan for the scientific illustration.

---

**Appendix 1.**
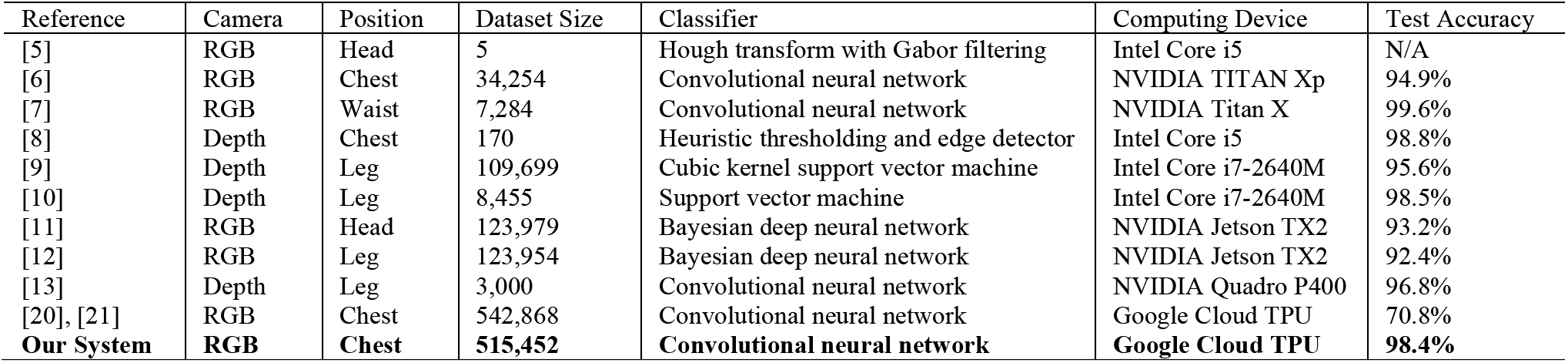
Summary of vision-based stair recognition systems for wearable robotics. Note that the dataset size (i.e., number of images) and test accuracy are only for the environment classes related to level-ground walking and stair ascent.

| Reference | Camera | Position | Dataset Size | Classifier | Computing Device | Test Accuracy |
| --- | --- | --- | --- | --- | --- | --- |
| [5] | RGB | Head | 5 | Hough transform with Gabor filtering | Intel Core i5 | N/A |
| [6] | RGB | Chest | 34,254 | Convolutional neural network | NVIDIA TITAN Xp | 94.9% |
| [7] | RGB | Waist | 7,284 | Convolutional neural network | NVIDIA Titan X | 99.6% |
| [8] | Depth | Chest | 170 | Heuristic thresholding and edge detector | Intel Core i5 | 98.8% |
| [9] | Depth | Leg | 109,699 | Cubic kernel support vector machine | Intel Core i7-2640M | 95.6% |
| [10] | Depth | Leg | 8,455 | Support vector machine | Intel Core i7-2640M | 98.5% |
| [11] | RGB | Head | 123,979 | Bayesian deep neural network | NVIDIA Jetson TX2 | 93.2% |
| [12] | RGB | Leg | 123,954 | Bayesian deep neural network | NVIDIA Jetson TX2 | 92.4% |
| [13] | Depth | Leg | 3,000 | Convolutional neural network | NVIDIA Quadro P400 | 96.8% |
| [20], [21] | RGB | Chest | 542,868 | Convolutional neural network | Google Cloud TPU | 70.8% |
| **Our System** | **RGB** | **Chest** | **515,452** | **Convolutional neural network** | **Google Cloud TPU** | **98.4%** |

## References

[1] M. R. Tucker et al., “Control strategies for active lower extremity prosthetics and orthotics: A review,” J. NeuroEngineering Rehabil., vol. 12, no. 1, p. 1, 2015.

[2] A. E. Patla, “Understanding the roles of vision in the control of human locomotion,” Gait Posture, vol. 5, no. 1, pp. 54–69, Feb. 1997.

[3] M. Grimmer, R. Riener, C. J. Walsh, and A. Seyfarth, “Mobility related physical and functional losses due to aging and disease – A motivation for lower limb exoskeletons,” J. NeuroEngineering Rehabil., vol. 16, no. 1, p. 2, Jan. 2019.

[4] A. J. Young and D. P. Ferris, “State of the art and future directions for lower limb robotic exoskeletons,” IEEE Trans. Neural Syst. Rehabil. Eng, vol. 25, no. 2, pp. 171–182, Feb. 2017.

[5] N. E. Krausz and L. J. Hargrove, “Recognition of ascending stairs from 2D images for control of powered lower limb prostheses,” in 2015 7th International IEEE/EMBS Conference on Neural Engineering (NER), Montpellier, France, Apr. 2015, pp. 615–618.

[6] B. Laschowski, W. McNally, A. Wong, and J. McPhee, “Preliminary design of an environment recognition system for controlling robotic lower-limb prostheses and exoskeletons,” in 2019 IEEE 16th International Conference on Rehabilitation Robotics (ICORR), Toronto, ON, Canada, Jun. 2019, pp. 868–873.

[7] G. Khademi and D. Simon, “Convolutional neural networks for environmentally aware locomotion mode recognition of lower-limb amputees,” in ASME Dynamic Systems and Control Conference (DSCC), Park City, Utah, USA, Oct. 2019, p. 11.

[8] N. E. Krausz, T. Lenzi, and L. J. Hargrove, “Depth sensing for improved control of lower limb prostheses,” IEEE Trans. Biomed. Eng., vol. 62, no. 11, pp. 2576–2587, Nov. 2015.

[9] Y. Massalin, M. Abdrakhmanova, and H. A. Varol, “User-independent intent recognition for lower limb prostheses using depth sensing,” IEEE Trans. Biomed. Eng., vol. 65, no. 8, pp. 1759–1770, Aug. 2018.

[10] H. A. Varol and Y. Massalin, “A feasibility study of depth image based intent recognition for lower limb prostheses,” in 2016 38th Annual International Conference of the IEEE Engineering in Medicine and Biology Society (EMBC), Orlando, FL, USA, Aug. 2016, pp. 5055–5058.

[11] B. Zhong, R. L. da Silva, M. Li, H. Huang, and E. Lobaton, “Environmental context prediction for lower limb prostheses with uncertainty quantification,” IEEE Trans. Autom. Sci. Eng., vol. 18, no. 2, pp. 458–470, Apr. 2021.

[12] B. Zhong, R. L. da Silva, M. Tran, H. Huang, and E. Lobaton, “Efficient environmental context prediction for lower limb prostheses,” IEEE Trans. Syst. Man Cybern. Syst., pp. 1–15, 2021.

[13] K. Zhang et al., “A subvision system for enhancing the environmental adaptability of the powered transfemoral prosthesis,” IEEE Trans. Cybern., vol. 51, no. 6, pp. 3285–3297, Jun. 2021.

[14] Y. LeCun, Y. Bengio, and G. Hinton, “Deep learning,” Nature, vol. 521, no. 7553, pp. 436–444, May 2015.

[15] B. Laschowski, W. McNally, A. Wong, and J. McPhee, “ExoNet database: Wearable camera images of human locomotion environments,” Front. Robot. AI, vol. 7, p. 562061, Dec. 2020.

[16] B. Laschowski, W. McNally, A. Wong, and J. McPhee, “Environment classification for robotic leg prostheses and exoskeletons using deep convolutional neural networks,” Front. Neurorobotics, vol. 15, p. 730965, Feb. 2022.

[17] A. G. Howard et al., “MobileNets: Efficient convolutional neural networks for mobile vision applications,” arXiv, Apr. 2017.

[18] M. Sandler, A. Howard, M. Zhu, A. Zhmoginov, and L.-C. Chen, “MobileNetV2: Inverted residuals and linear bottlenecks,” arXiv, Jan. 2018.

[19] M. Abadi et al., “TensorFlow: Large-scale machine learning on heterogeneous distributed systems,” arXiv, Mar. 2016.

[20] B. Laschowski, “Energy regeneration and environment sensing for robotic leg prostheses and exoskeletons,” PhD Thesis, University of Waterloo, 2021.

[21] B. Laschowski, W. McNally, A. Wong, and J. McPhee, “Computer vision and deep learning for environment-adaptive control of robotic lower-limb exoskeletons,” in 2021 43rd Annual International Conference of the IEEE Engineering in Medicine & Biology Society (EMBC), Mexico, Nov. 2021, pp. 4631–4635.

[22] D. P. Kingma and J. Ba, “Adam: A method for stochastic optimization,” arXiv, Dec. 2014.

[23] A. G. Kurbis, B. Laschowski, and A. Mihailidis, “StairNet: A computer vision dataset for stair recognition,” IEEE DataPort, Apr. 2022.

[24] J. Deng, W. Dong, R. Socher, L.-J. Li, K. Li, and L. Fei-Fei, “ImageNet: A large-scale hierarchical image database,” in 2009 IEEE Conference on Computer Vision and Pattern Recognition (CVPR), Miami, FL, USA, p. 8.

